# Modality-independent coding of scene categories in prefrontal cortex

**DOI:** 10.1101/142562

**Authors:** Yaelan Jung, Bart Larsen, Dirk B. Walther

**Affiliations:** 1Department of Psychology, University of Toronto.

**Author notes:** Correspondence should be addressed to Yaelan Jung, 100 St. George Street, Toronto, ON, M5S 3G3, Canada.

## Abstract

Natural environments convey information through multiple sensory modalities, all of which contribute to people’s percepts. Although it has been shown that visual or auditory content of scene categories can be decoded from brain activity, it remains unclear where and how humans integrate different sensory inputs and represent scene information beyond a specific sensory modality domain. To address this question, we investigated how categories of scene images and sounds are represented in several brain regions. A mixed gender group of healthy human subjects participated the present study, where their brain activity was measured with fMRI while viewing images or listening to sounds of different places. We found that both visual and auditory scene categories can be decoded not only from modality-specific areas, but also from several brain regions in the temporal, parietal, and prefrontal cortex. Intriguingly, only in the prefrontal cortex, but not in any other regions, categories of scene images and sounds appear to be represented in similar activation patterns, suggesting that scene representations in the prefrontal cortex are modality-independent. Furthermore, the error patterns of neural decoders indicate that category-specific neural activity patterns in the middle and superior frontal gyri are tightly linked to categorization behavior. Our findings demonstrate that complex scene information is represented at an abstract level in the prefrontal cortex, regardless of the sensory modality of the stimulus.

**Statement of Significance:** Our experience in daily life requires the integration of multiple sensory inputs such as images, sounds, or scents from the environment. Here, for the first time, we investigated where and how in the brain information about the natural environment from multiple senses is merged to form modality-independent representations of scene categories. We show direct decoding of scene categories across sensory modalities from patterns of neural activity in the prefrontal cortex. We also conclusively tie these neural representations to human categorization behavior based on the errors from the neural decoder and behavior. Our findings suggest that the prefrontal cortex is a central hub for integrating sensory information and computing modality-independent representations of scene categories.

## Introduction

Imagine taking a walk on the beach. Your sensory experience would include the sparkle of the sun’s reflection on the water, the sound of the crushing waves, and the smell of ocean air. Even though the brain has separate, clearly delineated processing channels for all of these sensory modalities, we still have the integral experience of being at the beach. What are the neural systems underlying this convergence of different sensory inputs, which allow us to form such a rich representation of our environment? Here we show that several brain regions contain neural representations of visual and auditory scene information with varying degrees of cross-modal integration.

Neural mechanisms underlying the perception of visual scenes has been studied extensively for the last two decades, showing a hierarchical structure from posterior to anterior parts of visual cortex with increasing level of abstraction. Starting from low-level features, such as orientation, represented in primary visual cortex, the level of representation becomes more abstract, through intermediate-level features in the occipital place area (Dilks, Julian, Paunov, & Kanwisher, 2013; MacEvoy & Epstein, 2007), to the representation of local scene geometry and scene category in the parahippocampal place area (Epstein & Kanwisher, 1998 Walther, Caddigan, Fei-Fei, & Beck, 2009) and the embedding of a specific scene into real-world topography in the RSC and hippocampus (Morgan *et al*., 2011). Does this abstraction continue beyond the visual domain? To identify representations of scene contents beyond this visual processing hierarchy, we here investigated neural representation of scene contents delivered from visual and auditory cues.

Previous work has identified neural representations that are not constrained to a particular sense for other types of stimuli. Several brain areas have been shown to integrate signals from more than one sense (Calvert, 2001; Driver & Noesselt, 2008), such as posterior superior temporal sulcus (Beauchamp *et al*., 2004), the posterior parietal cortex (Cohen and Anderson, 2004; Molholm *et al*., 2006; Sereno and Huang, 2006) and the prefrontal cortex (Sugihara et al, 2006; Romanski, 2007). Some of these areas show similar neural activity patterns when the same information is delivered from different senses for various stimuli, such as objects (Man, Kaplan, Damasio & Meyer, 2012), emotions (Müller, Cieslik, Turetsky, & Eickhoff, 2012), or face/voice identities (Park *et al*., 2010). Despite these observations, little is known about how *scene* information is processed beyond the sensory modality domain.

In real world settings, our perception of scenes typically relies on multiple senses. Therefore, we postulate that there should exist a stage of modality-independent representation of scenes beyond the visual hierarchy, which should generalize information across different modality channels. We hypothesized that PFC may play a role in representing scene categories beyond the modality domain based on the previous research showing that PFC shows categorical representations of visual information (Freedman, Riesenhuber, Poggio, & Miller, 2001; Miller, Freedman, & Wallis, 2002; Walther *et al*., 2009).

The present study investigates modality-independent scene representation using MVPA of fMRI data. First, we identified brain areas that process both visual and auditory scene information by decoding neural representations of scene categories elicited by scene images and sounds separately. We hypothesized that some of these areas, which contain both visual and auditory scene content, might not compute modality-independent representations but process each modality input separately. To test this idea, we performed cross-decoding analysis between the two modalities to establish cross-modal areas. Furthermore, we examined whether representations of scene categories in one modality is degraded by conflicting stimulation in the other domain (i.e. a beach image and an office sound), which shows whether a brain area integrate information from different sensory domains.

Finally, to tie neural representations of scene contents to human categorization behavior, we compared the errors of neural decoders to those from a separate behavioral experiment. Among the multisensory brain regions that we investigated, only prefrontal regions contain modality-independent representations of scene categories, integrate information from different modality channels and reflect human categorization behavior.

## MATERIALS AND METHODS

We posited four different models of how visual and auditory information can be processed within a brain region, a purely visual model, a purely auditory model, a multi-modal model with separate but intermixed neural populations for processing visual and auditory information, and a cross-model model with neurons truly integrating visual and auditory information (Fig 1C). Experimental conditions and analysis protocols were designed to discriminate between these models (Fig 1A, B).

**Figure 1.**
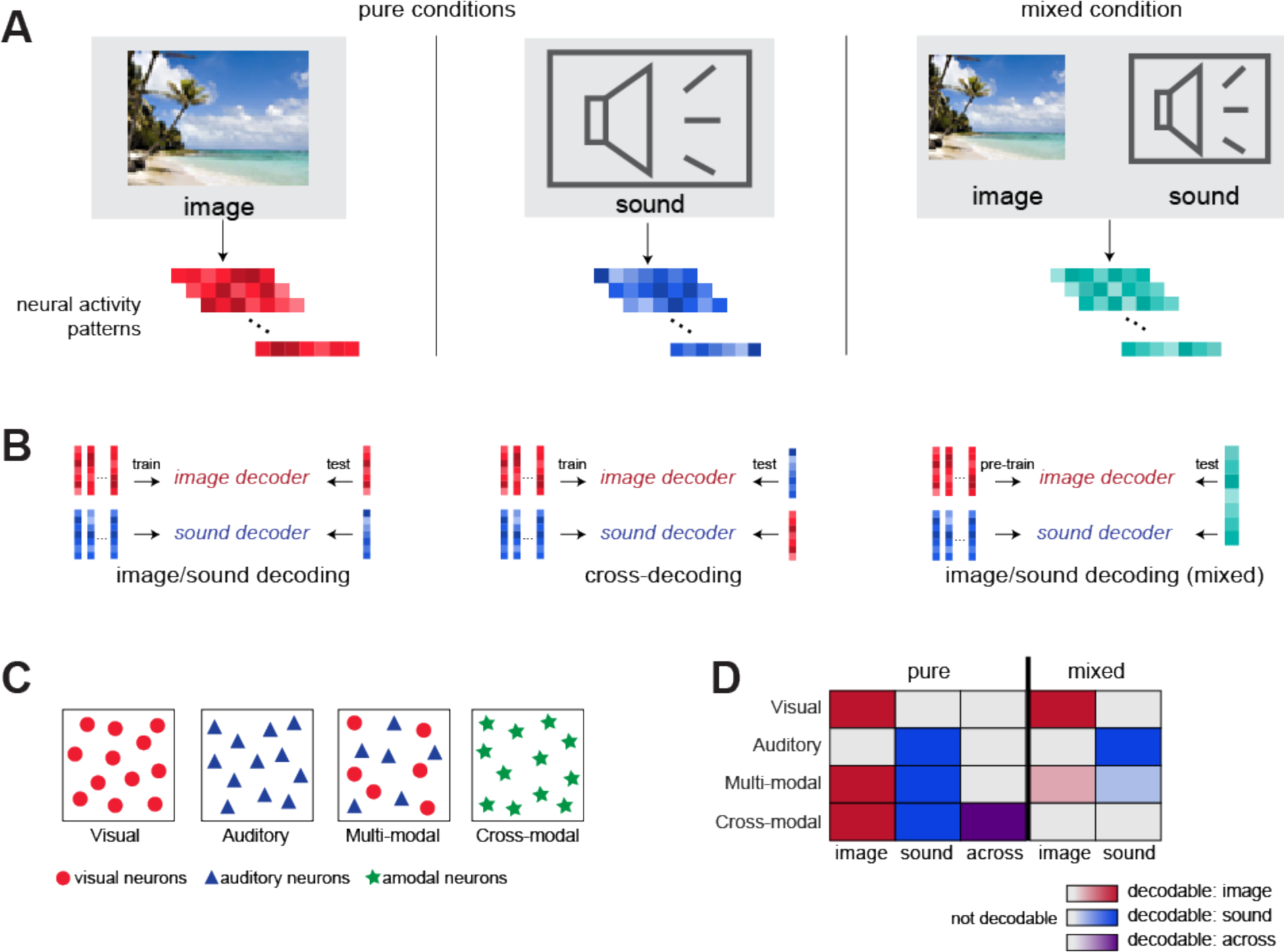
**(A)** Illustration of the image, sound, and mixed conditions. In the pure image and sound conditions, either images or sounds from four different scene categories were presented while neural activity patterns of participants were recorded. In the mixed condition, both images and sounds were presented simultaneously, but they were always from different categories (i.e. a beach images with city sounds). **(B)** Multivariate analysis of fMRI data for decoding image and sound category, cross-decoding across image and sound, and decoding image and sound from the mixed condition. **(C)** Models for separate brain areas dedicated to visual, auditory, multi-modal, and cross-modal processing. **(D)** Predictions for decoding performance of each model in different conditions (full color: decodable, grey: not decodable)

Figure 1D shows predicted results for each of the four models. Specifically, we expect that primary visual and auditory regions will contain neurons dedicated to the processing their respective modality exclusively. In these regions, scene categories should be decodable from the corresponding modality condition only, but not across modalities. Also, conflicting information from the other modality should not interfere with the neural representation from the preferred modality. In multi-modal regions, both visual and auditory information should be processed in anatomically collocated but functionally separate neural populations. Therefore, we expect that both image and sound categories can be decoded, but decoding across modalities should not be possible. Conflicting information from the other modality should not interfere with image and sound processing, as the information from each modality channel is processed separately. It is conceivable that these multimodal areas give precedence to one particular sense, not processing both equally, when there is conflicting information from each channel. Finally, scene categories from both modalities should be decodable from cross-modal regions, as long as they are consistent. Hence, both image and sound categories should be decodable, and scene category decoding should generalize across modalities. However, conflicting information from one modality will interfere with processing of information from the other modality in this region, so that decoding of scene categories will be degraded when the information from images and sounds is inconsistent.

### Participants

Thirteen subjects (18 to 25 years old, 6 females; 7 males) participated in the fMRI experiment. All participants were in good health with no past history of psychiatric or neurological disorders and reported having normal hearing and normal or corrected-to-normal vision. They gave written informed consent before the experiment began according to the Institutional Review Board of The Ohio State University.

A separate group of 25 undergraduate students from the University of Toronto (18 to 21 years old, 16 females; 9 males) participated in the behavioral experiment for course credit. All participants had normal hearing and normal or corrected-to-normal vision and gave written informed consent. The experiment was approved by the Research Ethics Review Board of the University of Toronto.

### Stimuli

In the fMRI experiment, 640 color photographs of different scene categories (beaches, forests, cities, & offices) were used. The images have previously been rated as the best exemplars of their categories from a data base of about 4000 images that were downloaded from the internet (Torralbo *et al*. 2013). Images were presented at a resolution of 800×600 pixels using a Christie DS+6K-M projector operating at a refresh rate 60Hz. Images subtended approximately 21×17 degrees of visual angle.

Sixty-four sound clips representing the same four scene categories (beaches, forests, cities, or offices) were used as auditory stimuli. The sound clips were purchased from various commercial sound libraries and edited to 15 seconds of length. Because of this relatively longer presentation time for each audio exemplar, fewer exemplars were used compared to those in the image condition. Perceived loudness was equated using Replay Gain as implemented in the Audacity sound editing software (Audacity Team, 2012). In a pilot experiment, the sound clips were correctly identified and rated as highly typical for their categories by 14 naïve subjects.

The same visual and the auditory stimuli were used in the behavioral experiment. In the visual part of the experiment, 400 images were used for practice blocks (key-category association and staircase-wise practice), and the 240 images were used in the main testing blocks. Images were presented on a CRT screen at a resolution of 800×600 pixels and subtended approximately 29 x 22 degrees of visual angle. The resolution of the monitor was 1024 x 768 with a refresh rate at 150 Hz. Images were followed by a perceptual mask, which was generated by synthesizing a mixture of textures reflecting all four scene categories (Portilla & Simoncelli, 2000).

### Procedure and Experiment design

#### fMRI experiment

The fMRI experiment consisted of three conditions; the image condition, the sound condition, and the mixed condition, in which both images and sounds presented concurrently.Participants’ brains were scanned during twelve experimental runs, four runs for each condition. Each run started with the instruction asking participants to attend, for the duration of the run, to either images (image runs and half of the mixed runs) or sounds (sound runs and the other half of the mixed runs).

Runs contained eight blocks, two for each scene category, interleaved with 12.5 s fixation periods to allow for the hemodynamic response to return to baseline levels. The beginning and the end of a run also included a fixation period of 12.5 sec. The order of blocks within runs and the order of runs were counter-balanced across participants. Mixed runs were only presented after at least two pure image and sound runs. Stimuli were arranged into eight blocks of 15 seconds duration. During image blocks participants were shown ten color photographs of the same scene category for 1.5 seconds each. During sound blocks they were shown a blank screen with a fixation cross, and a 15-second sound clip was played using Sensimetrics S14 MR-compatible in-ear noise-canceling headphones at approximately 70 dB. During mixed blocks participants were shown images and played a sound clip of a different scene category at the same time. A fixation cross was presented throughout each block, and subjects were instructed to maintain fixation. Each run lasted 3 min and 52.5 seconds.

#### fMRI data acquisition and preprocessing

Imaging data were recorded on a 3 Tesla Siemens MAGNETOM Trio MRI scanner with a 12-channel head coil at the Center for Cognitive and Behavioral Brain Imaging (CCBBI) at The Ohio State University. High resolution anatomical images were acquired with a 3D-MPRAGE (magnetization-prepared rapid acquisition with gradient echo) sequence with sagittal slices covering the whole brain; inversion time = 930ms, repetition time (TR) = 1900ms, echo time (TE) = 4.68ms, flip angle = 9°, voxel size = 1 x 1 x 1 mm, matrix size = 224 x 256 x 160 mm. Functional images for the main experiment were recorded with a gradient echo, echo-planar imaging sequence with a volume repetition time (TR) of 2.5 s, an echo time (TE) of 28ms and a flip angle of 78 degrees. 48 axial slices with 3 mm thickness were recorded without gap, resulting in an isotropic voxel size of 3 x 3 x 3 mm.

FMRI data were motion corrected to one EPI image (the 72nd volume of the 10th run), followed by spatial smoothing with a Gaussian kernel with 2 mm full width at half maximum (FWHM) and temporal filtering with a high-pass filter at 1/400 Hz. Data were normalized to percent signal change by subtracting the mean of the first fixation period in each run and dividing by the mean across all runs. The effects of head motion (6 motion parameters) and scanner drift (second degree polynomial) were regressed out using a general linear model (GLM). The residuals of this GLM analysis were averaged over the duration of individual blocks, resulting in 96 brain volumes that were used as input for a multi-voxel pattern analysis (MVPA). Preprocessing was performed using AFNI (Cox, 1996).

#### Behavioral experiment

The behavioral experiment consisted of two parts, a visual and an auditory part. The order of the two parts was randomized for each participant.

The visual part consisted of two phases; practice and testing. Participants performed two types of practice; one for the key-category association and the other for the fast image presentation. During the first block of practice, photographs of natural scenes were presented for 200ms (stimulus onset asynchrony, SOA), immediately followed by a perceptual mask for 500ms. Participants were asked to press one of four keys (‘a’, ‘s’, ‘k’, ‘l’) within 2 second of stimulus onset. They received acoustic feedback (a short beep) when they made an error. Assignment of the keys to the four scene categories (beaches, forests, cities, offices) was randomized for each participant. After participants achieved 90% accuracy in this key practice phase, they completed four additional practice blocks (40 trials each), during which the SOA was linearly decreased to 26.7ms. The main testing phase consisted of six blocks (40 trials each) of 4AFC scene categorization with a fixed SOA of 26.7ms and without feedback.

In the auditory part, participants listened to scene soundscapes of 15 seconds in length. They indicated their categorization decision by pressing one of four keys (same key assignment as in the visual part). In order to make the task difficulty comparable to the image categorization task, we overlaid pure-tone noise onto the original sound clips. Noise consisted of 30ms snippets of pure tones, whose frequency was randomly chosen between 50 and 2000 Hz with 3 ms of fade-in and fade-out (linear ramp). Based on a pilot experiment, we set the relative volume of the noise stimulus to nine times the volume of the scene sounds. To familiarize participants with the task, they first performed 4 practice trials. Participants were asked to respond with the key corresponding to the correct category, starting from 8 seconds after the onset of the sound and without an upper time limit. Participants were encouraged not to deliberate on the answer but to respond as quickly and as accurately as possible.

### Data analysis and Statistical analysis

#### Defining Regions of Interest

Regions of interest in visual cortex were defined using functional localizer scans, which were performed at the end of the same session as the main experiment. Retinotopic areas in early visual cortex were identified using the meridian stimulation method (Kastner, Weerd,Desimone, & Ungerleider, 1998). The vertical and horizontal meridians of the visual field were stimulated alternatingly with wedges with flickering checkerboard pattern. Boundaries between visual areas were outlined on the computationally flattened cortical surface. The boundary between V1 and V2 was identified as the first vertical meridian activity, the boundary between V2 and V3 as the first horizontal meridian, and the boundary between V3 and V4 (lower bank of the calcarine fissure only) as the second vertical meridian. To establish the anterior boundary of V4, we stimulated the upper and lower visual field in alternation with flickering checkerboard patterns. The anterior boundary of V4 was found by ensuring that both upper and lower visual field are represented in V4. Participants maintained central fixation during the localizer scan.

To define high-level visual areas, we presented participants with blocks of images of faces, scenes, objects, and scrambled images of objects. FMRI data from this localizer were pre-processed the same way as the main experiment data, but spatially smoothed with a 4 mm FWHM Gaussian filter. Data were further processed using a general linear model (3dDeconvolve in Afni) with regressors for all four image types. ROIs were defined as contiguous clusters of voxels with significant contrast (q < 0.05, corrected using false discovery rate) of: scenes > (faces and objects) for PPA, RSC (Epstein and Kanwisher, 1998) and OPA (Dilks *et al*., 2013); and objects > (scrambled objects) for LOC (Malach *et al*., 1995). Voxels that could not be uniquely assigned to one of the functional ROIs were excluded from the analysis.

Anatomically defined ROIs were extracted using a probability atlas in AFNI (DD Desai MPM; Destrieux *et al*., 2010): middle temporal gyrus (MTG), superior temporal gyrus (STG), superior temporal sulcus (STS), angular gyrus (AG), superior parietal gyrus (SPG), intraparietal sulcus (IPS), middle frontal gyrus (MFG), superior frontal gyrus (SFG), and inferior frontal gyrus (IFG) with pars opercularis, pars orbitalis, and pars triangularis. Anatomical masks for primary auditory cortex and its subdivisions (Planum Temporale, Posteromedial Heschl’s gyrus, Middle Heschl’s gyrus, Anterolateral Heschl’s gyrus, & Planum Polare) were made available by Sam Norman-Haignere (Norman-Haignere, Kanwisher, & McDermott, 2013). After nonlinear alignment of each participants’ anatomical image to MNI space using Afni’s 3dQwarp function, the inverse of the alignment was used to project anatomical ROI masks back into original subject space using 3dNwarpApply. All decoding analyses, including for the anatomically defined ROIs, were performed in original subject space.

#### Multi-voxel pattern analysis

For each participant, we trained a linear support vector machine (SVM; using LIBSVM, Chang & Lin, 2001) to assign the correct scene category labels to the voxel activations inside an ROI based on the fMRI data from all runs except one. The SVM decoder then produced predictions for the labels of the data in the left-out run. This leave-one-run-out (LORO) cross validation procedure was repeated with each run being left out in turn, thus producing predicted scene category labels for all runs. Decoding accuracy was assessed as the fraction of blocks with correct category labels. Group-level statistics was computed over all thirteen participants using one-tailed t tests, determining if decoding accuracy was significantly above chance level (0.25).Significance of the t-test was adjusted for multiple comparisons using false discovery rate (FDR) (Westfall & Young, 1993).

To curb over-fitting of the classifier to the training data, we reduced the dimensionality of the neural data by selecting a subset of voxels in each ROI. Voxel selection was performed by ranking voxels in the training data according to the F statistics of a one-way ANOVA of each voxel’s activity with scene category as the main factor (Pereira, Mitchell, & Botvinick. 2009). We determined the optimal number of voxels by cross validation within the training data. In the case of cross validation analysis of the pure image and sound conditions, voxel selection was performed in a nested cross-validation, using the training data of each cross validation fold. Optimal voxel numbers varied by ROI and participant but were generally between 100 and 1000 (mean voxel number averaged across all ROIs and subjects = 107.125).

#### Error correlations

Category label predictions of the decoder were recorded in a confusion matrix, whose rows indicate the category of the stimulus, and whose columns represent the category predictions by the decoder. Diagonal elements indicate correct predictions, and off-diagonal elements represent decoding errors. Neural representations of scene categories were compared with human behavior by correlating the error patterns (the off-diagonal elements of the confusion matrices) between neural decoding and behavioral responses (Walther, Beck, & Fei-Fei 2012). Statistical significance of the correlations was established non-parametrically against the null distribution of all error correlations that were obtained by jointly permuting the rows and columns of one of the confusion matrices in question (24 possible permutations of four labels). Error correlations were considered as significant when none of the correlations in the null set exceeded the correlation for the correct ordering of category labels (p < 0.0417).

To assess the similarity between neural representations and the physical characteristics of the stimuli, we constructed simple computational models of scene categorization based on low-level stimulus features. Scene images were filtered with a bank of Gabor filters with four different orientations at four scales. Images were categorized based on the resulting feature vector in a 16-fold cross validation, using a linear SVM, resulting in a classification accuracy of 85.8% (chance: 25%).

Physical properties of the sounds were assessed using a cochleagram, which mimics the biomechanics of the human ear (Meddis, Hewitt, & Shackleton, 1990; Wang & Brown, 2006). The cochleagrams of individual sound clips were integrated over their duration and subsampled to 128 frequency bands, resulting in a biomechanically realistic frequency spectrum. The activation of the frequency bands was used as input to a linear SVM, which predicted scene categories of sounds in a 16-fold cross validation. The classifier accurately categorized 57.8% of the scene sounds (chance: 25%). Error patterns from computational analysis for images and sounds were correlated with those obtained from the neural decoder.

Error patterns of human observers were obtained from the behavioral experiment. Mean accuracy was 76.6% for the visual task (std = 12.35%, mean RT = 885.6 ms) and 78.1% for the auditory task (std = 6.86%, mean RT = 7.84 sec). Behavioral errors were recorded in confusion matrices, separately for images and sounds. CM rows indicate the true category of the stimulus, and columns indicate participants’ responses. Individual cells contain the relative frequency of the responses indicated by the columns to stimuli indicated by the rows. Group-mean CMs were compared to confusion matrices derived from neural decoding.

#### Searchlight analysis

To explore representations of scene categories outside of pre-defined ROIs, we performed a searchlight analysis. We defined a cubic “searchlight” of 7×7×7 voxels (21×21×21 mm). The searchlight was centered on each voxel in turn (Kriegeskorte, Göbel, & Bandettini, 2006), and LORO cross-validation analysis was performed within each searchlight location using a Gaussian Naïve Bayes classifier until all voxels served as the center of the searchlight (Searchmight Toolbox; Pereira & Botvinick 2011). Decoding accuracy as well as the full confusion matrix at a given searchlight location, were assigned to the central voxel.

For group-analysis, we first co-registered each participant’s anatomical brain to the Montreal Neurological Institute (MNI) 152 template using a diffeomorphic transformation as calculated by AFNI’s 3dQWarp. We then used the same transformation parameters to register individual decoding accuracy maps to MNI space using 3dNWarpApply, followed by spatial smoothing with a 4 mm FWHM Gaussian filter. To identify voxels with decodable categorical information at the group level, we performed one-tailed t-tests to test whether decoding accuracy at each searchlight location was above chance (0.25). After thresholding at p < .01 (one-tailed) we conducted a cluster-level correction for multiple comparisons, applying a minimum cluster size of 15 voxels, the average cluster size obtained from the α probability simulations conducted for each participant.

## RESULTS

### Decoding scene categories of images and sounds

To assess neural representations of scene categories from images and sounds, we performed multivoxel pattern analysis for each ROI. A linear support vector machine (SVM; using LIBSVM, Chang & Lin, 2001) was trained using neural activity patterns with category labels and then tested to determine if a trained classifier can predict the scene category in a leave-one-run-out cross validation.

Figure 2 illustrates decoding accuracy in each condition for various brain areas (see Table 1 for the results of statistical analysis). As shown in previous studies (Choo & Walther, 2016; Park, Brady, Greene, & Oliva, 2011; Walther *et al*., 2009; Walther, Chai, Caddigan, Beck, & Fei-Fei, 2011; Kravitz, Peng, & Baker, 2011), both early visual areas V1 through V4 and high-level visual areas, including the parahippocampal place area (PPA), the retrosplenial cortex (RSC), the occipital place area (OPA), and lateral occipital complex (LOC), show category-specific scene representations. We were also able to decode the scene categories from activity elicited by sounds of the respective natural environments in primary auditory cortex (A1) as well as its anatomical sub-divisions (Fig. 2).

**Figure 2.**
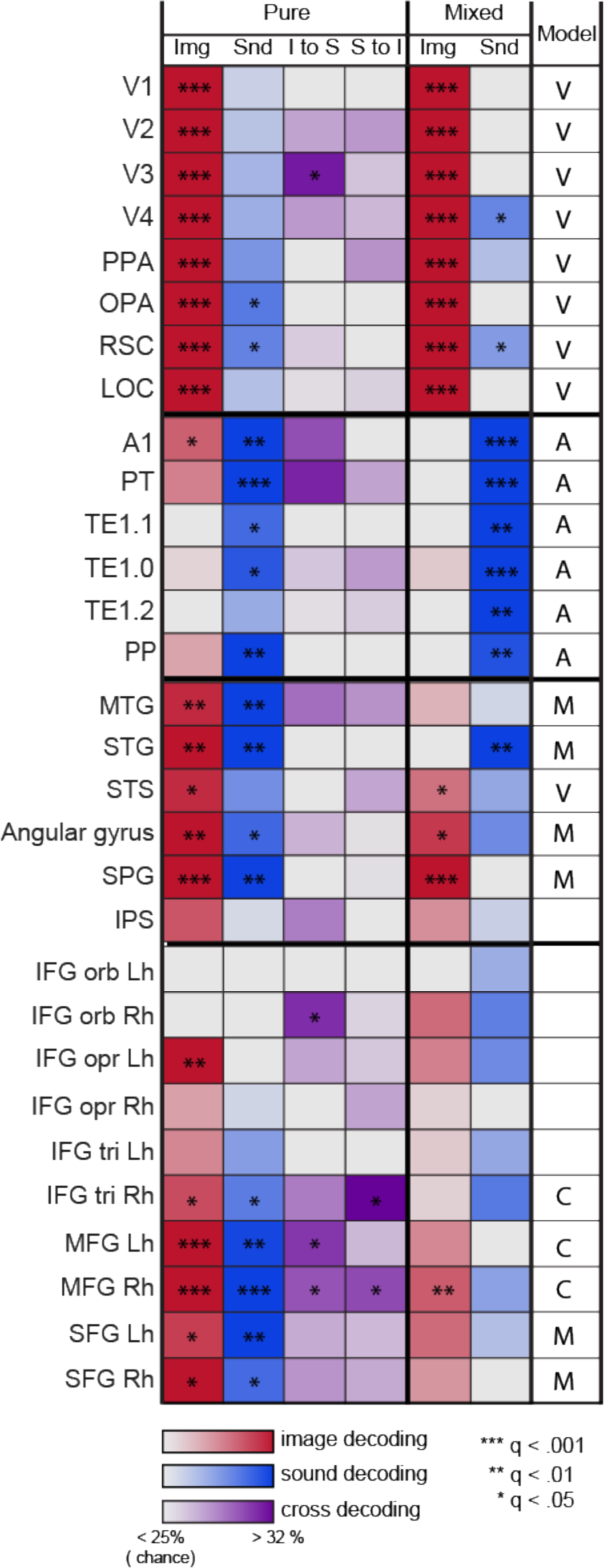
Decoding accuracy in each ROI (listed in each row) for each condition (listed in each column) are illustrated with the degree of color saturation. Decoding accuracies for the image scene categories are illustrated in red, those for the sound categories are illustrated in blue, and those for the cross decoding analysis are illustrated in purple. The first four columns are results from the image and sound conditions and the two columns on the right are the results from the mixed condition. The last column indicates the corresponding models for each ROI, whose predictions are comparable to the results in each ROI; V: visual, A: auditory, M: multi-modal, C: cross-modal. Significance of the one-sample t-tests (one-tailed) was adjusted for multiple comparisons using false discovery rate, *q < .05, **q < .01, ***q < .001.

**Table 1.**
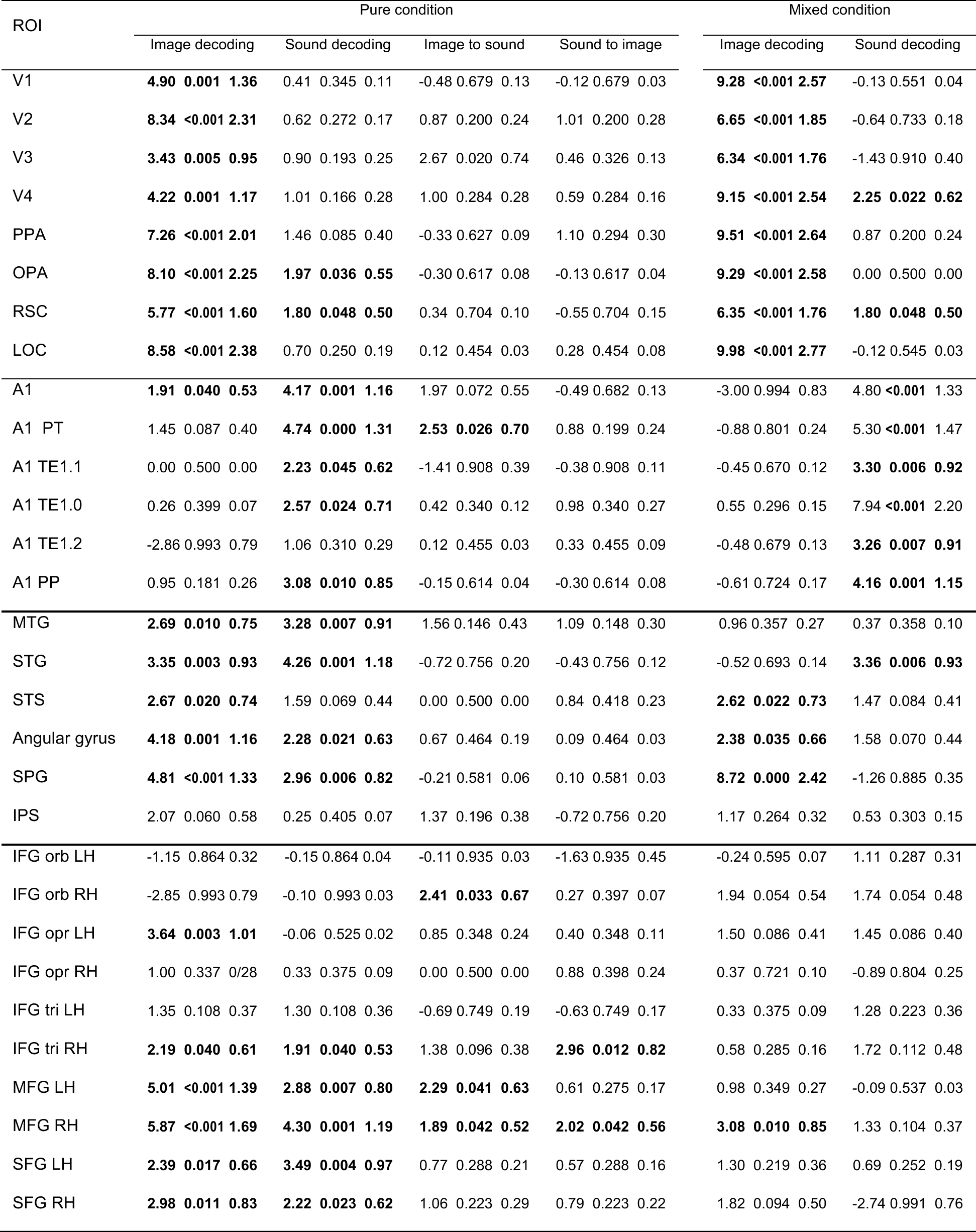
Statistical results of decoding performance in each ROI for each type of decoding analysis: t-value, q value (p corrected for multiple comparison), Cohen’s d. One-sample t-test (one tailed) was performed to see whether neural decoder’s performance is over the chance (25%) and the significance of the test was adjusted using false discovery rate (FDR) ; the degree of freedom in each t-test was 12, and the significant tests *(q* < .05) are highlighted in bold font.

Unlike previous reports showing that auditory content can be decoded from early visual cortex (Paton, Petro, & Muckli, 2016; Vetter *et al*., 2014), we did not find representations of auditory scene information in V1 to V4. However, we were able to decode auditory scene categories in higher visual areas, the OPA (30.5%, t(12) = 1.966, *q* = 0.036, *d* = 1.36) and the RSC (31.3%; t(12) = 1.803, *q* =0.048, *d* = 0.5). Intriguingly, we could also decode scene categories from images in A1 with a decoding accuracy of 29.8%, t(12) = 1.910, *q* = 1.91, *d* = 0.53.

Having found modality-specific representations of scene categories in visual and auditory cortices, we aimed to identify scene representations in areas which are not limited to a specific sensory modality. We could decode categories of both visual and auditory scenes in several temporal and parietal regions (Fig. 2): the middle temporal gyrus (MTG), the superior temporal gyrus (STG), the superior parietal gyrus (SPG), and the angular gyrus (AG). In the superior temporal sulcus (STS), we could decode scene categories only from the images, not from the sounds. Although previous studies have suggested that the intraparietal sulcus (IPS) is involved in audiovisual processing (Calvert, Hansen, Iversen, & Brammer, 2001), we could not decode visual or auditory scene categories in the IPS.

Next, we examined whether the prefrontal cortex (PFC) also showed category-specific representations for both visual and auditory scene information. Previous studies have found strong hemispheric specialization in PFC (Gaffan & Harrison,1991; Goel, *et al*., 2007; Slotnick & Moo, 2006). We therefore analyzed functional activity in PFC separately by hemisphere. We were able to decode visual scene categories significantly above chance from the left inferior frontal gyrus (IFG), pars opercularis, the right IFG, pars triangularis, and in both hemispheres from the middle frontal gyrus (MFG) and the superior frontal gyrus (SFG; Fig. 2 & Table 1). The categories of scene sounds were decodable in the right IFG, pars triangularis, as well as the MFG and SFG in both hemispheres (Fig. 2 & Table 1).

Although the temporal, parietal, and prefrontal cortex all showed both visual and auditory scene representations, this does not necessarily imply that these areas process scene information beyond the sensory modality domain. Neural representation of scene categories in the cross-modal regions should not merely consist of co-existing populations of neurons with visually and auditorily triggered activation patterns (see Fig 1c, *the multi-modal model);* the voxels in these ROIs should be activated equally by visual and auditory inputs if they represent the same category. In other words, if the neural activity pattern elicited by watching a picture of a forest reflects the scene category of *forest,* then this neural representation should be similar to that elicited by listening to forest sounds (see Fig 1c, *the cross-modal model).* We aimed to explicitly examine whether scene category information in the prefrontal areas transcends sensory modalities using cross-decoding analysis between the image and sound conditions.

### Cross-modal decoding

For the cross-decoding analysis, we trained the decoder using the image labels from the image runs and then tested whether it could correctly predict the categories of scenes presented as sounds in the sound runs. We also performed the reverse analysis, training the decoder on the sound runs and testing it on the image runs (Fig. 2).

Cross-decoding from images to sounds succeeded in the MFG in both hemispheres and in the right IFG, pars orbitalis. The right MFG and the right IFG, pars triangularis, showed significant decoding accuracy for cross-decoding from sounds to images (Fig. 2). However, cross-decoding was not possible in either direction anywhere in sensory cortices or temporal and parietal cortices which have significant decoding of both image and sound categories. Although V3 and the planum temporale (PT) shows significant decoding from the cross-decoding analysis of images to sounds (see Fig 2 and Table 1), it is hard to interpret these findings as equivalent to those in prefrontal regions since they only show significant decoding of either image or sound categories in the straight decoding analysis. These results therefore suggest that only prefrontal areas contain modality-independent representations of scene categories with similar neural activity patterns from visual and auditory scene information.

### Presenting images and sounds concurrently

As a further test to examine the cross-modal nature of scene category representations in PFC, we used an interference condition, in which we presented images and sounds from incongruous categories simultaneously. If a population of neurons encodes a scene category independently of sensory modality, then we should see a degradation of the category representation in the presence of a competing signal from the other modality. If, on the other hand, two separate but intermixed populations of neurons encode the visual and auditory categories, respectively, then we should be able to still decode the category from at least one of the two senses.

To decode scene categories from this mixed condition, we created an image and a sound decoder by training separate classifiers with data from the image-only and the sound-only conditions. We then tested these decoders with neural activity patterns from the mixed condition, using either image or sound labels as ground truth (Fig. 2). As the training and the test data are from separate sets of runs, cross-validation was not needed for this analysis.

We were able to decode visual and auditory scene categories from the respective sensory brain areas, even in the presence of conflicting information from the other modality. In temporal and parietal ROIs, we could decode scene categories, but these ROIs were no longer multimodal; they only represented scene categories in either the visual or auditory domain but no longer both (Fig 2 & Table 1). These findings suggest that these ROIs contain separate but intermixed neural populations for visual and auditory information. For ROIs in PFC, we found that conflicting audiovisual stimuli interfered heavily with representations of scene categories (Fig. 2). Scene categories could no longer be decoded in PFC from either modality, except for visual scenes in the right MFG. Presumably, this break-down of the decoding of scene categories is due to the conflicting information from the two sensory modalities arriving at the same cross-modal populations of neurons.

### Error correlations

To further explore the characteristics of the neural representations of scene categories, we compared the patterns of errors from the neural decoders with those from human behavior as well as with the physical attributes of the stimuli. If neural representation of scenes in a certain brain region is used directly for categorical decisions, then error patterns from this ROI should be similar to errors made in behavioral categorization (Walther *et al*., 2009). However, in early stages of neural processing, scene representations might reflect the physical properties of the scene images or sounds. In this case, the error patterns of the decoders should resemble the errors that a classifier would make solely based on low-level physical properties.

To assess similarity of representations, we correlated the patterns of errors (off-diagonal elements of the confusion matrices, see Methods) between the neural decoders, physical structure of the stimuli, and human behavior. Statistical significance of the correlations was established with non-parametric permutation tests. Here we considered error correlations to be significant when none of the correlations in the null set exceeded the correlation of the correct ordering of the categories *(p* < 0.0417).

Behavioral errors from image categorization were not correlated with the errors derived from image properties (*r* = − .458, *p* = .917), suggesting that behavioral judgment of scene categories was not directly driven by low-level physical differences between the images. There was, however, a positive error correlation between the auditory task and physical properties of sounds (*r* = .407, *p* < .0417).

For the image condition, errors from neural decoders were similar to those from image structure in early visual cortex and similar to human behavioral errors in high-level visual areas (Fig 3A); in early visual cortex, decoding errors were positively correlated with image structure (V1: *r* = .746, *p* <.0417; V2: *r* = .451, *p* = .083) but not with behavioral errors (V1: *r* = −0.272, *p*= .0708; V2: *r* = −.132, *p* = .667). V3 and V4 showed no strong correlation with stimulus structure (V3: *r* = −0.171, *p* = .708; V4: *r* = .076, *p* = .667) or behavior (V3: *r* = .356, *p* = .125; V4: *r* = .250, *p*= .125). In high-level scene-selective areas, decoding errors were positively correlated with image behavior (PPA: *r* = .570, *p* < .0417; RSC: *r* = 0.637, *p* < .0417; not in OPA: *r* = .183, *p*= .292), but not with the error patterns representing image structure (PPA: *r* = .0838, *p* = 0.333; RSC: *r* = −.230, *p* = 0.708; OPA: *r* = .099, *p* = .458).

**Figure 3.**
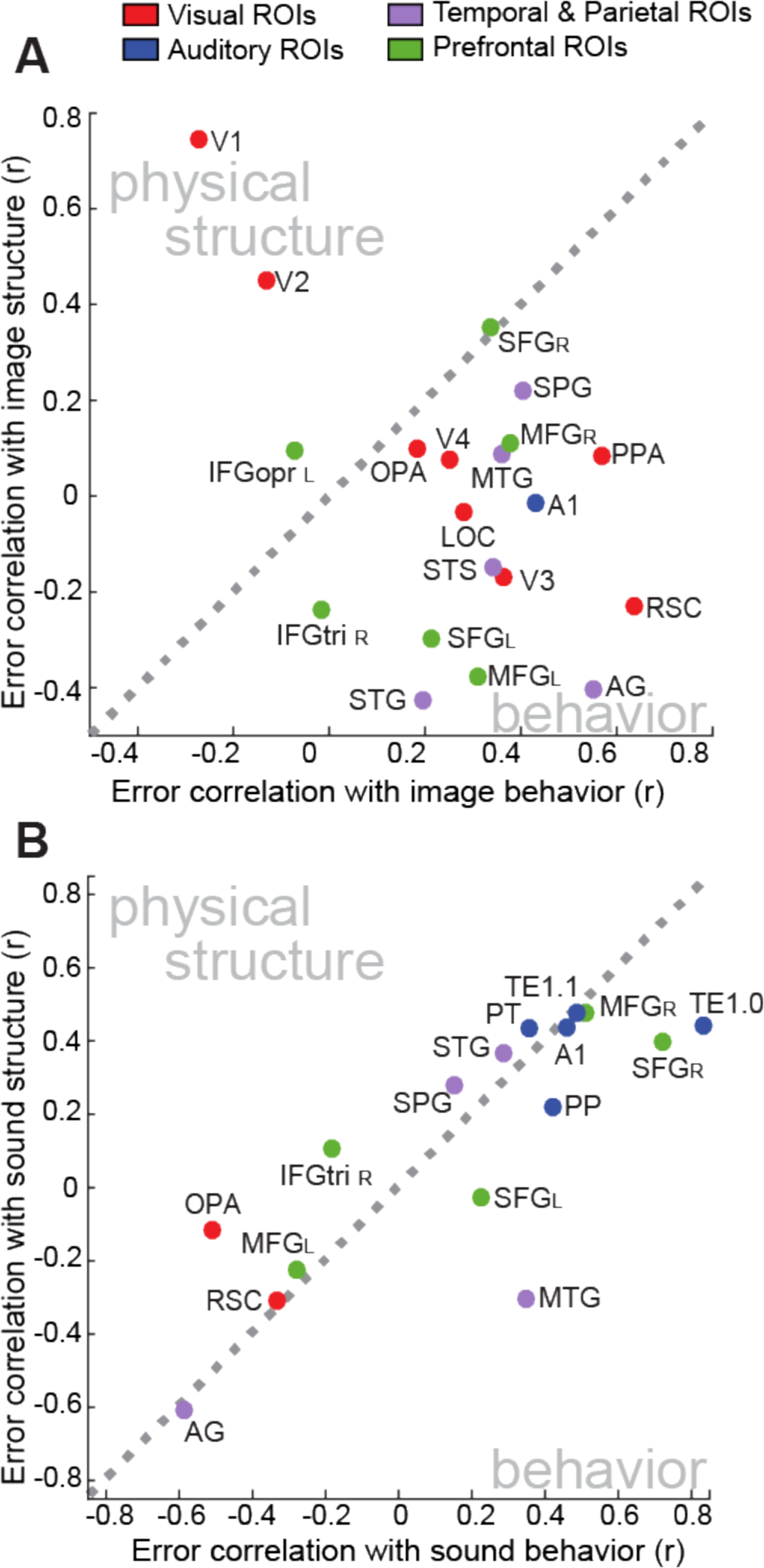
Error correlations of the neural decoder with behavior and stimuli structure in the image **(A)** and sound **(B)** conditions. Diagonal axes indicate the points where the error correlations with behavior and with stimuli structure are equal. In the image condition, we see a clear progression from V1 through V2-4 to higher-level visual areas (in red), moving from strong error correlation with image structure to strong error correlation with visual behavior. Error patterns from decoding image categories from PFC (in green) are most similar to visual behavior. In the sound condition, all ROIs are close to the diagonal, because sound structure and sound behavior error patterns are significantly correlated with each other. We see A1 and its subdivisions (in blue) high along the diagonal, indicating strong similarity with both sound structure and sound behavior. OPA and RSC (in red) show low error correlation with either structure or behavior. Prefrontal ROIs (in green) are distributed along the diagonal, with right MFG and right SFG showing high and left MFG showing low error correlations.

Errors from the neural decoders in SPG were positively correlated with image behavior (r = .404, *p* = < .0417) but not with image structure (r = .220, *p* = .167). The errors from MTG, STS, and AG also showed high correlation with image behavior errors, but did not reach significance in the permutation test (MTG: *r* = .360, *p* = .083; STS: *r* = .348, *p* = .083; AG: *r* = .552, *p* = .083; Fig. 3A). Finally, in PFC, errors from the image decoders in the right MFG and SFG show significant correlation with image behavior (right MFG: *r* = .377, *p* < .0417; right SFG, *r* = .338, *p* < .0417). The left hemisphere of these ROIs also showed positive correlations but not significantly (left MFG: *r* = .309, *p* = .083; left SFG: *r* = .212, *p* = .208). On the other hand, the left IFG, pars opercularis, and the right IFG, pars triangularis, showed no error correlation at all with either image behavior or structure (Fig. 3A).

In the sound condition, error patterns from sound structure as well as sound behavior were positively correlated with decoding errors from A1, even though the permutation test did not reach significant level (with sound structure: *r* = .438, *p* = .083; with sound behavior: *r* = .46; *p*= .125). However, three of the five anatomical sub-divisions of Heschl’s Gyrus showed significant correlation with sound structure (TE1.1: *r* = .478, *p* < .0417; TE1.0: *r* = .443, *p* <.0417). TE1.0 also showed significant error correlations with human behavior for sounds (r = .83, *p* < .0417; Fig. 3B).

None of the temporal or parietal ROIs showed significant error correlations of decoding sounds with sound structure or behavior (Fig. 3B). In PFC, we found that errors from the right SFG significantly correlate with both sound structure (r = .397, *p* < .0417) and sound behavior (r = .721, *p* < .0417). The left SFG and the right MFG also showed high correlations with sound behavior but not significantly (left SFG: *r* = .224, *p* = .208; right MFG: *r* = .486, *p* = .125; Fig. 3b).

### Whole-brain analysis

In order to explore representations of scene categories beyond the pre-defined ROIs, we performed a whole-brain searchlight analysis with a size of 7×7×7 voxels (21×21×21 mm) cubic searchlight. The same LORO cross-validation analysis for image and sound conditions as well as the same two cross-decoding analyses as for the ROI-based analysis were performed at each searchlight location, followed by a cluster-level correction for multiple comparisons. For each decoding condition, we found several spatial clusters with significant decoding accuracy. Some of these clusters confirmed the pre-defined ROIs, others revealed scene representations in unexpected regions beyond the ROIs.

For the decoding of the image categories, we found a large cluster of 15359 voxels with significant decoding accuracy for decoding scene images, spanning most of visual cortex. In accordance with the ROI-based analysis, we also found clusters in prefrontal cortex, overlapping with the right middle frontal gyrus (MFG), and the superior frontal gyrus (SFG), the inferior frontal gyrus (IFG) and MFG in the left hemisphere. See Figure 4A and Table 2 for a complete list of clusters. Decoding of sound categories produced two large clusters in the two hemispheres, which overlap with primary auditory cortices, as well as clusters near the right insula and in left STG (see Fig. 4A and Table 2).

**Figure 4.**
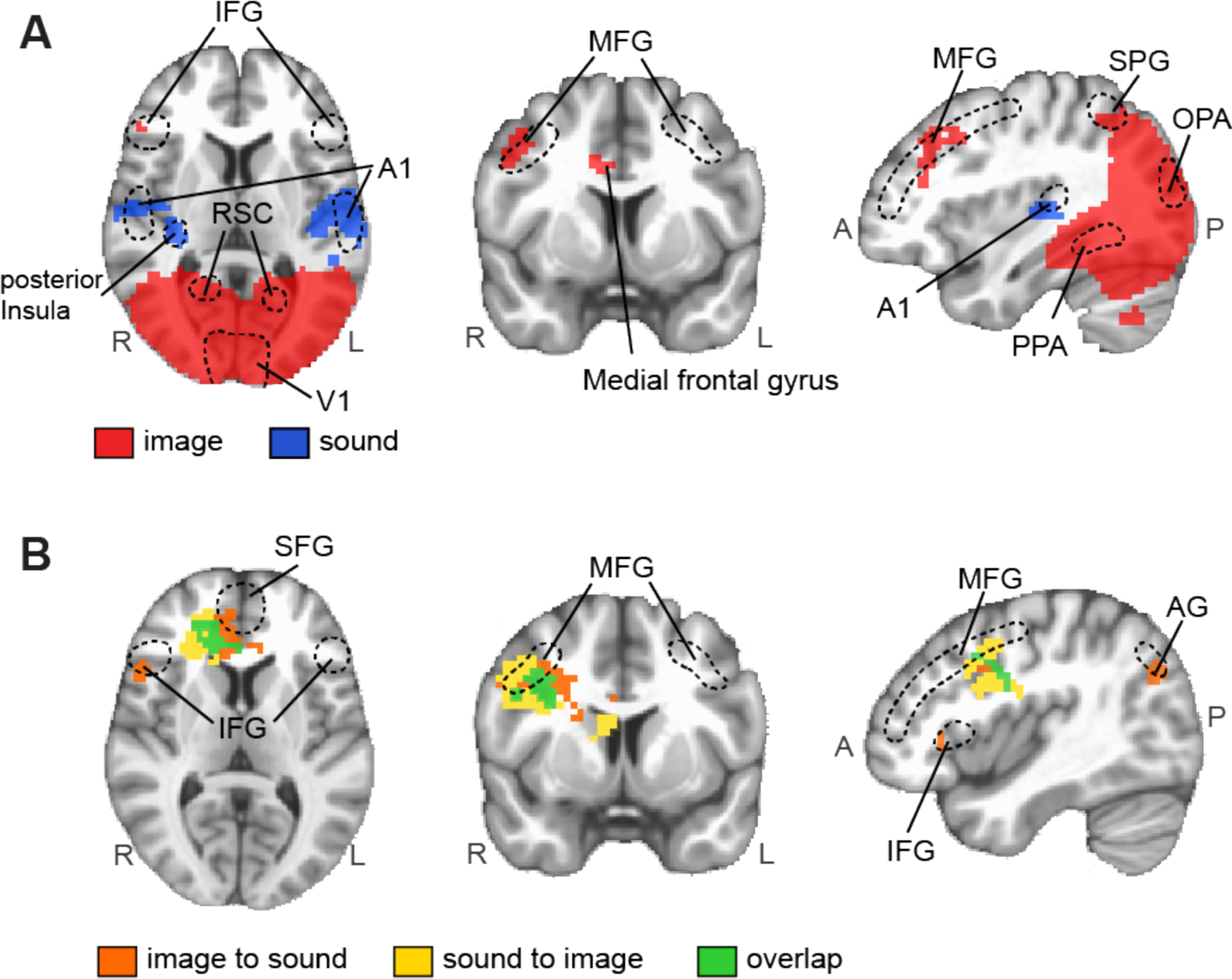
Searchlight maps for (A) decoding image and sound categories and (B) cross-decoding across image and sound categories (thresholded at p < .01). **(A)** Searchlight locations for image decoding overlap with the parahippocampal place area (PPA), the retrosplenial cortex (RSC), the occipital place area (OPA), the superior parietal gyrus (SPG), the inferior frontal gyrus (IFG), the middle frontal gyrus (MFG), and the superior frontal gyrus (SFG), which are highlighted in red. Searchlight locations for sound decoding overlap with A1, which are highlighted in blue. **(B)** Searchlight locations for cross-decoding analysis (image to sound, sound to image, or both) overlap with PPA, IFG, MFG, and SFG, which are highlighted in orange, yellow, or green respectively.

**Table 2.** Clusters identified in the searchlight analysis for decoding of image and 786 sound scene categories (thresholded at p < .01).

| Decoding Condition | Peak (MNI coordinates) | | | | Volume (μl) | Description |
| --- | --- | --- | --- | --- | --- | --- |
|  | x | y | z | Accuracy (%) |  |  |
| Image | -1.5 | 81.9 | 7 | 40.1 | 407262 | Middle occipital gyri, parahippocampal gyri, precuneus, cuneus, superior parietal lobule, right angular gyrus |
|  | -46.1 | -10.2 | 34 | 30.8 | 7212 | Right middle frontal gyrus |
|  | -4.5 | -40 | 31 | 29.6 | 4216 | Right medial frontal gyrus |
|  | 40 | 1.6 | 31 | 28.8 | 1273 | Left Inferior frontal gyrus |
|  | 34.2 | -48.9 | -8 | 28.0 | 1087 | Left middle frontal gyrus |
|  | 13.4 | -60.8 | 25 | 28.6 | 663 | Left superior frontal gyrus |
|  | 19.3 | -48.9 | -14 | 27.5 | 424 | Left superior frontal gyrus |
| Sound | 52 | 13.5 | 4 | 31.5 | 13364 | Left superior temporal gyrus |
|  | -52 | 13.5 | 7 | 31.1 | 2439 | Right superior temporal gyrus |
|  | -37.2 | 19.5 | 7 | 29.2 | 1193 | Right posterior insula |

Even though we were able to find ROIs that allowed for decoding of both images and sounds, we could not find any searchlight locations where this was possible. This may be due to spatial smoothing introduced in the alignment to the standard brain and due the different classifier - we used a Gaussian Naïve Bayesian classifier for the searchlight analysis and a linear SVM for the ROI-based analysis.

We found several significant clusters in the right prefrontal cortex that allowed for cross-decoding between images and sounds (Fig. 4B). The image-to-sound condition produced clusters with significant decoding accuracy in the right MFG, SFG, and IFG as well as right superior temporal gyrus (STG), right angular gyrus (AG), right inferior temporal gyrus (ITG), and left middle occipital gyrus. The sound-to-image condition resulted in clusters in the right MFG, the IFG and the anterior cingulate gyrus as well as the right pre/postcentral gyri and the parahippocampal gyrus (See Fig 4B and Table 3 for details). We found two compact clusters that allowed for cross-decoding in both directions in the right prefrontal cortex, overlapping with the right IFG (124 voxels) and MFG (208 voxels). Cross-decoding between images and sounds was not possible anywhere else.

**Table 3.** Clusters identified in the searchlight analysis for cross-decoding of scene images and sound (thresholded at p < .01).

| Decoding Condition | Peak (MNI coordinates) | | | | Volume (μl) | Description |
| --- | --- | --- | --- | --- | --- | --- |
|  | x | y | z | Accuracy (%) |  |  |
| Image To Sound | -10.4 | -31.1 | 16 | 29.2 | 18800 | Right anterior cingulate, right medial frontal gyrus, right superior frontal gyrus |
|  | -31.2 | -10.2 | 40 | 28.1 | 9731 | Right middle frontal gyrus, right inferior frontal gyrus |
|  | -55 | 61.1 | 13 | 28.8 | 2811 | Right superior temporal gyrus, right angular gyrus, right middle temporal gyrus |
|  | -31.2 | -34 | 22 | 27.6 | 1273 | Right middle frontal gyrus, right inferior frontal gyrus |
|  | 28.2 | 22.5 | 22 | 27.3 | 955 | Left insula |
|  | 37.2 | 16.5 | 31 | 27.3 | 610 | Left postcentral gyrus, left precentral gyrus |
|  | -52 | -22.1 | 7 | 28.0 | 504 | Right inferior frontal gyrus |
|  | 16.4 | 87.9 | 13 | 26.9 | 451 | Left middle occipital gyrus, left cuneus |
|  | 22.3 | -31.1 | 10 | 27.3 | 398 | Left anterior cingulate |
| Sound To Image | -22.3 | -37 | 10 | 28.1 | 10527 | Right medial frontal gyrus, right anterior cingulate, right inferior frontal gyrus |
|  | -49.1 | -10.2 | 40 | 28.8 | 8777 | Right middle frontal gyrus, right inferior frontal gyrus, right insula |
|  | 31.2 | 16.5 | 28 | 27.8 | 5197 | Left postcentral gyrus, left precentral gyrus, left insula |
|  | -34.2 | 22.5 | -8 | 27.0 | 1299 | Right parahippocampal gyrus, right hippocampus |

We compared error patterns from the neural decoders to stimulus properties and human behavior for all searchlight locations that allowed for decoding of scene categories (Fig. 5 & Table 4). Clusters in visual cortex, overlapping with V1-V4, showed significant error correlations with image properties. Error patterns from searchlight locations in the superior parietal gyrus (SPG) and the parahippocampal gyrus, overlapping with the PPA, correlated with errors from human behavior for image categorization. In general, we observed a posterior-to-anterior (PA) trend, with voxels in the posterior (low-level) visual regions more closely matched to stimulus properties and with voxels more anterior (high-level) visual regions more closely related to behavior.

**Figure 5.**
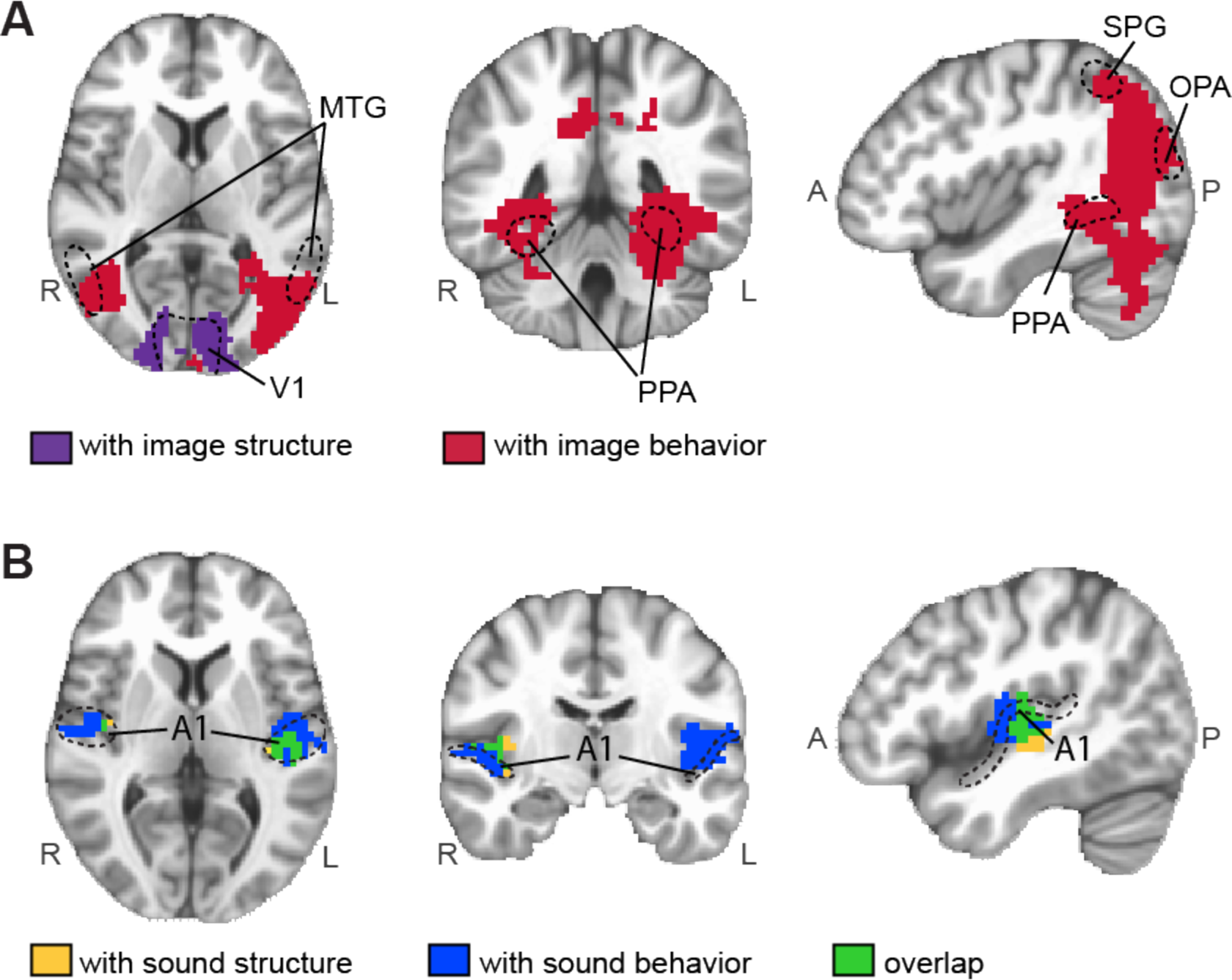
Searchlight maps or error correlations in the image **(A)** and sound **(B)** condition (thresholded at p < .01). Note that **(B)** uses a more compact range of axial slices than **(A)**. There were no significant error correlations outside of this range of slices. **(A)** Searchlight locations with significant error correlations with image structure overlap with V1 and V2 (highlighted in purple). Searchlight locations with significant error correlations with image behaviors overlaps with the parahippocampal place area (PPA), and the retrosplenial cortex (RSC). **(B)** Searchlight locations with significant error correlations with sound structure, and sound behaviors (or both) overlap with A1 and its subdivisions such as the planum temporale (PT), posteromedial Heschl’s gyrus (TE1.1), middle Heschl’s gyrus (TE1.0), anterolateral Heschl'’s gyrus (TE1.2), and the planum polare (PP).

**Table 4.** Clusters identified in the searchlight analysis for error correlations in the image and the sound conditions (thresholded at p < .05).

| Error correlation | | Peak (MNI coordinates) | | | | Volume (μl) | Description | ROIs with overlap |
| --- | --- | --- | --- | --- | --- | --- | --- | --- |
|  |  | x | y | z | correlation coefficient |  |  |  |
| image condition | with image structure | 16.4 | 96.8 | -2 | .868 | 36354 | middle occipital gyrus, inferior occipital gyrus, lingual gyrus, cuneus | V1, V2, V3, V4 |
|  | with image behavior | 1.5 | 99.8 | 16 | .829 | 172753 | middle temporal gyrus, middle occipital gyrus, superior occipital gyrus, inferior parietal lobule, superior parietal lobule | PPA, OPA, RSC, LOC |
| sound condition | with sound structure | 43.1 | 19.5 | 1 | .648 | 3182 | left superior temporal gyrus | left A1 |
|  |  | -49.1 | 10.6 | 7 | .701 | 530 | right precentral gyrus, right superior temporal gyrus | right A1 |
|  | with sound behavior | 52 | 10.6 | 10 | .757 | 7318 | left superior temporal gyrus | left A1 |
|  |  | -55 | 7.6 | 7 | 0.787 | 1671 | right precentral gyrus, right superior temporal gyrus | right A1 |

Clusters in bilateral STG, overlapping with A1, showed significant error correlations with sound properties and behavioral errors for sound categorization. Within this cluster, we see the same posterior-to-anterior (PA) trend, with posterior voxels being more closely related to sound properties and more anterior voxels being more closely related to behavioral categorization of scene sounds (Fig. 5B).

## DISCUSSION

The present study investigated where and how scene information from different sensory domains forms modality-independent representations of scene categories. We have found that both visual and auditory stimuli of the natural environment elicit representations of scene categories in sub-regions of prefrontal cortex (PFC). These neural representations of scene categories generalize across sensory modalities and resemble human categorization behavior, suggesting that scene representations in PFC reflect scene categories not constrained to a specific sensory domain. To our knowledge, our study is the first to demonstrate a neural representation of scenes at such an abstract level.

Three distinct characteristics support the idea that neural representations of scene categories in PFC are distinct from those in modality-specific areas such as the visual or the auditory cortices or other multisensory areas. First, both image and sound categories could be decoded from the same areas in PFC. Thus, it can be inferred that neural representations of scene categories in PFC are not limited to a specific sensory modality channel. Second, the representations in PFC could be cross-decoded from one modality to the other, showing that the category-specific neural activity patterns were similar across the sensory modalities. Third, when subjects were presented with incongruous visual and auditory scene information simultaneously, it was no longer possible to decode scene categories in PFC, whereas modality-specific areas as well as multimodal areas still carried the category-specific neural activity patterns. This result shows that inconsistent information entering through the two sensory channels in the mixed condition interferes, preventing the formation of c of scene categories in PFC.

Although scene categories could be decoded from both images and sounds in several ROIs in the temporal and parietal lobes, cross-decoding across sensory modalities was not possible there, suggesting that neural representations elicited by visual inputs were not similar to those elicited by auditory inputs. Further supporting the idea that visual and auditory representations are separate but intermixed in these regions, decoding of scene categories from the visual or auditory domain was still possible in the presence of a conflicting signal in the other domain. These findings suggest that even though information from both visual and auditory inputs is present in these regions (Beauchamp *et al*., 2004; Calvert *et al*., 2001), scene information is computed separately for each sensory modality, unlike in PFC. The discrimination between multi-modal and truly cross-modal representations is not possible with the univariate analysis techniques used in those studies.

Analysis of decoding errors demonstrated that the category representations in the visual areas have a hierarchical organization. In the early stage of processing, categorical representations are formed based on the physical properties of visual inputs, whereas in the later stage, the errors of neural decoders correlate with human behavior, confirming previous findings which mainly focused on the scene-selective areas (Walther *et al*., 2009; 2011; Choo & Walther,2016). Significant error correlation between human behavior and the neural decoders in prefrontal areas confirms that this hierarchical organization is extended to PFC, beyond the modality-specific areas PPA, OPA, and RSC.

Intriguingly, no similar hierarchical structure of category representations was found in the auditory domain. Both types of errors, the errors representing the physical properties and those from human behavior, were correlated to the errors of neural decoder in the A1. This difference between the visual and the auditory domain might reflect the fact that much of auditory processing occurs in sub-cortical regions, before the information arrives in auditory cortex. Thus, if the auditory scene processing is relying on a neural architecture with a hierarchy, it might not be easily detectable with fMRI. A recent study by Teng, Sommer,Pantazis and Oliva (2017) showed evidence suggesting a potential structure under auditory scene processing, by finding that different types of auditory features in a scene, reverberant space and source identity, are processed at a different time course. Further investigation with other various recording techniques, such as MEG/EEG in combination with fMRI, as well as with computational modeling (Cichy & Teng, 2016) necessitates for a better understanding of the neural mechanism of auditory scene processing.

Previous fMRI studies have shown that auditory content can be decoded from early visual cortex, suggesting cross-modal interactions in the modality-specific areas (Paton *et al*., 2016; Vetter *et al*., 2014). Although we did not observe representations of auditory scenes in the early visual areas, our data show that auditory content can be decoded from high-level scene-selective areas (RSC and OPA). Visual content can be decoded from A1. This seeming inconsistency with previous studies might be driven by different levels of complexity and diversity of the auditory stimuli used in each experiment: Vetter *et al*. (2014) used few exemplars of object-level sounds. We used more complex sounds with multiple objects overlapping in time, recorded from real-world settings, whose statistics could only be classified at the category level. Thus, category-specific visual imagery caused by auditory stimulation may underlie the decoding of auditory scene categories in RSC and OPA. These findings lead to a host of further questions for future research, such as how these visual and auditory areas are functionally connected, whether the multisensory areas mediate this interaction between the visual and auditory areas by sending feedback signals, or whether these cross-modal representations can influence or interfere with perceptual sensitivity in each sensory domain.

The whole-brain searchlight analysis confirmed the findings of our ROI-based analysis. In the image and sound decoding analyses, we found clusters with significant decoding accuracies in the visual and the auditory areas as well as in the temporal, the parietal, and the prefrontal regions. Furthermore, the clusters in the prefrontal areas showed significant accuracy in the cross-decoding analysis, while the clusters in other modality-specific or multimodal areas did not, supporting the view that only representations in the prefrontal cortex transcend sensory modalities. In the analysis of decoding errors, we observed that the errors of the image decoders were significantly correlated with human categorization behavior in scene-selective areas PPA and RSC as well as in the superior parietal gyrus, consistent with previous work by our group (Walther *et al*., 2009; 2011; Choo & Walther, 2016).

Previous studies addressing the integration of audiovisual information to form a modality-independent representations have employed univariate analysis (Beauchamp *et al*., 2004; Downar *et al*., 2000) or correlations of content-specific visual and auditory information in the brain (Hsier *et al*., 2012). These methods do not distinguish between co-activation from multiple senses and modality-independent processing. Recent studies using MVPA have shown that visual and auditory information about objects (Man *et al*., 2012) or emotions (Peelen, Atkinson & Vuilleumier, 2010) evokes similar neural activity patterns across different senses, suggesting that stimulus content is represented independently of sensory modality at later stages of sensory processing. Unlike the present study, however, these studies report that areas in temporal or parietal cortex are involved in this multimodal integration. One reason for this difference could be that real-world scenes are more variable in their detailed sensory representation, typically including multiple visual and auditory cues. Furthermore, we here consider representations of scene *categories* as opposed to object *identity* (Man *et al*., 2012). Our results indicate that generalization across sensory modalities at the level of scene categories occurs only in PFC. The same brain regions have been found to be involved in purely visual categorization and category learning (Freedman *et al*., 2001; Mack, Preston, & Love,2013; Meyers, Freedman, Kreiman, Miller, & Poggio, 2008; Miller & Cohen, 2001; Wood & Grafman, 2003).

In a recent review, Grill-Spector and Weiner (2014) suggested that the ventral temporal cortex contains a hierarchical structure for visual categorization, which has the more exemplar-specific representations in posterior areas, but the more abstract representations in anterior areas of the ventral temporal cortex. In the present study, we show that the posterior-to-anterior hierarchy of levels of abstraction extends to the PFC, which represents scene categories beyond the sensory modality domain. The abstraction and generalization across sensory modalities is likely to contribute to the efficiency of cognition by representing similar concepts in a consistent manner, even when the physical signal might be delivered via different sensory channels (Huth, de Heer, Griffiths, Theunissen, & Gallant, 2016).

## Acknowledgements

We thank Michael Mack and Heeyoung Choo for their helpful comments on the early version of this manuscript. This work is supported by NSERC Discovery Grant (#498390) and Canadian Foundation for Innovation (#32896).

## Notes

Conflict of interest: The authors declare no conflict of interest.

## References

Beauchamp, M., Lee, K., Argall, B., & Martin, A. (2004). Integration of auditory and visual information about objects in superior temporal sulcus. Neuron, 41, 809–823.

Calvert, G. A. (2001). Crossmodal processing in the human brain: insights from functional neuroimaging studies. Cerebral Cortex, 11(12), 1110–1123.

Calvert, G. A., Hansen, P. C., Iversen, S. D., & Brammer, M. J. (2001). Detection of audio-visual integration sites in humans by application of electrophysiological criteria to the BOLD effect. NeuroImage, 14(2), 427–38. http://doi.org/10.1006/nimg.2001.0812

Chang, C. C., & Lin, C. J. (2011). LIBSVM: a library for support vector machines. ACM Transactions on Intelligent Systems and Technology (TIST), 2(3), 27.

Choo, H., & Walther, D. B. (2016). Contour junctions underlie neural representations of scene categories in high-level human visual cortex. NeuroImage, 135, 32–44.

Cichy, R. M., & Teng, S. (2016). Resolving the neural dynamics of visual and auditory scene processing in the human brain: a methodological approach. Philos.Trans. R. Soc. B, 372, 1–11. http://doi.org/10.1098/rstb.2016.0108

Cohen, Y. E., & Andersen, R. A. (2004). Multimodal spatial representations in the primate parietal lobe. Crossmodal space and crossmodal attention, 99–121.

Cox, R. W. (1996). AFNI: software for analysis and visualization of functional magnetic resonance neuroimages. Computers and Biomedical research, 29(3), 162–173.

Destrieux, C., Fischl, B., Dale, A., & Halgren, E. (2010). Automatic parcellation of human cortical gyri and sulci using standard anatomical nomenclature. NeuroImage, 53(1), 1–15.

Dilks, D. D., Julian, J. B., Paunov, A. M., & Kanwisher, N. (2013). The occipital place area is causally and selectively involved in scene perception. Journal of Neuroscience, 33(4), 1331–1336.

Downar, J., Crowley, A. P., Mikulis, D. J., & Davis, K. D. (2000). A multimodal cortical network for the detection of changes in the sensory environment. Nature Neuroscience, 3(3), 277–283.

Driver, J., & Noesselt, T. (2008). Multisensory interplay reveals crossmodal influences on ‘sensory-specific’ brain regions, neural responses, and judgments. Neuron, 57(1), 11–23.

Epstein, R., & Kanwisher, N. (1998). A cortical representation of the local visual environment. Nature, 392(6676), 598–601.

Freedman, D. J., Riesenhuber, M., Poggio, T., & Miller, E. K. (2001). Categorical representation of visual stimuli in the primate prefrontal cortex. Science, 291(January), 312–317.

Gaffan, D., & Harrison, S. (1991). Auditory-visual associations, hemispheric specialization and temporal-frontal interaction in the rhesus monkey. Brain, 114(5), 2133–2144.

Goel, V., Tierney, M., Sheesley, L., Bartolo, A., Vartanian, O., & Grafman, J. (2007). Hemispheric specialization in human prefrontal cortex for resolving certain and uncertain inferences. Cerebral cortex, 17(10), 2245–2250

Grill-Spector, K., & Weiner, K. S. (2014). The functional architecture of the ventral temporal cortex and its role in categorization. Nature Reviews Neuroscience, 15(8), 536–548.

Hsieh, P.-J., Colas, J. T., & Kanwisher, N. (2012). Spatial pattern of BOLD fMRI activation reveals cross-modal information in auditory cortex. Journal of Neurophysiology, 107(12), 3428–3432.

Huth, A. G., de Heer, W. A., Griffiths, T. L., Theunissen, F. E., & Gallant, J. L. (2016). Natural speech reveals the semantic maps that tile human cerebral cortex. Nature, 532(7600), 453–458.

Kastner, S., De Weerd, P., Desimone, R., & Ungerleider, L. G. (1998). Mechanisms of directed attention in the human extrastriate cortex as revealed by functional MRI. Science, 282(5386), 108–111.

Kravitz, D. J., Peng, C. S., & Baker, C. I. (2011). Real-world scene representations in high-level visual cortex: it’s the spaces more than the places. Journal of Neuroscience, 31(20), 7322–7333.

Kriegeskorte, N., Goebel, R., & Bandettini, P. (2006). Information-based functional brain mapping. Proceedings of the National academy of Sciences of the United States of America, 103(10), 3863–3868.

MacEvoy, S. P., & Epstein, R. A. (2007). Position selectivity in scene-and object-responsive occipitotemporal regions. Journal of Neurophysiology, 98(4), 2089–2098.

Mack, M. L., Preston, A. R., & Love, B. C. (2013). Decoding the brain’s algorithm for categorization from its neural implementation. Current Biology, 23(20), 2023–2027.

Malach, R., Reppas, J. B., Benson, R. R., Kwong, K. K., Jiang, H., Kennedy, W. A., … & Tootell, R. B. (1995). Object-related activity revealed by functional magnetic resonance imaging in human occipital cortex. Proceedings of the National Academy of Sciences, 92(18), 8135–8139.

Man, K., Kaplan, J. T., Damasio, A., & Meyer, K. (2012). Sight and sound converge to form modality-invariant representations in temporoparietal cortex. Journal of Neuroscience, 32(47), 16629–16636.

Meddis, R., Hewitt, M. J., & Shackleton, T. M. (1990). Implementation details of a computation model of the inner hair-cell auditory-nerve synapse. The Journal of the Acoustical Society of America, 87(4), 1813–1816.

Meyers, E. M., Freedman, D. J., Kreiman, G., Miller, E. K., & Poggio, T. (2008). Dynamic population coding of category information in inferior temporal and prefrontal cortex. Journal of Neurophysiology, 100(June 2008), 1407–1419. http://doi.org/10.1152/jn.90248.2008

Miller, E. K., & Cohen, J. D. (2001). An integrative theory of prefrontal cortex function. Annual Review of Neuroscience, 24, 167–202. http://doi.org/10.1146/annurev.neuro.24.1.67

Miller, E. K., Freedman, D. J., & Wallis, J. D. (2002). The prefrontal cortex: categories, concepts and cognition. Philosophical Transactions of the Royal Society B: Biological Sciences, 357(1424), 1123–1136. http://doi.org/10.1098/rstb.2002.1099

Molholm, S., Sehatpour, P., Mehta, A. D., Shpaner, M., Gomez-Ramirez, M., Ortigue, S., … & Foxe, J. J. (2006). Audio-visual multisensory integration in superior parietal lobule revealed by human intracranial recordings. Journal of Neurophysiology, 96(2), 721–729.

Morgan, L. K., MacEvoy, S. P., Aguirre, G. K., & Epstein, R. A. (2011). Distances between real-world locations are represented in the human hippocampus. Journal of Neuroscience, 31(4), 1238–1245.

Muller, V. I., Cieslik, E. C., Turetsky, B. I., & Eickhoff, S. B. (2012). Crossmodal interactions in audiovisual emotion processing. NeuroImage, 60(1), 553–561.

Norman-Haignere, S., Kanwisher, N., & McDermott, J. H. (2013). Cortical pitch regions in humans respond primarily to resolved harmonics and are located in specific tonotopic regions of anterior auditory cortex. Journal of Neuroscience, 33(50), 19451–19469.

Park, S., Brady, T. F., Greene, M. R., & Oliva, A. (2011). Disentangling scene content from spatial boundary: complementary roles for the parahippocampal place area and lateral occipital complex in representing real-world scenes. Journal of Neuroscience, 31(4), 1333–1340.

Park, J. Y., Gu, B. M., Kang, D. H., Shin, Y. W., Choi, C. H., Lee, J. M., & Kwon, J. S. (2010). Integration of cross-modal emotional information in the human brain: an fMRI study. Cortex, 46(2), 161–169.

Paton, A., Petro, L., & Muckli, L. (2016). An Investigation of Sound Content in Early Visual Areas. Journal of Vision, 16(12), 153–153.

Peelen, M. V., Atkinson, A. P., & Vuilleumier, P. (2010). Supramodal representations of perceived emotions in the human brain. Journal of Neuroscience, 30(30), 10127–10134.

Pereira, F., & Botvinick, M. (2011). Information mapping with pattern classifiers: a comparative study. NeuroImage, 56(2), 476–496.

Pereira, F., Mitchell, T., & Botvinick, M. (2009). Machine learning classifiers and fMRI: a tutorial overview. NeuroImage, 45(1), S199–S209.

Portilla, J., & Simoncelli, E. P. (2000). A parametric texture model based on joint statistics of complex wavelet coefficients. International Journal of Computer Vision, 40(1), 49–70.

Romanski, L. M. (2007). Representation and integration of auditory and visual stimuli in the primate ventral lateral prefrontal cortex. Cerebral Cortex, 17(suppl_1), i61–i69.

Sereno, M. I., & Huang, R. S. (2006). A human parietal face area contains aligned head-centered visual and tactile maps. Nature Neuroscience, 9(10), 1337–1343.

Slotnick, S. D., & Moo, L. R. (2006). Prefrontal cortex hemispheric specialization for categorical and coordinate visual spatial memory. Neuropsychologia, 44(9), 1560–1568.

Sugihara, T., Diltz, M. D., Averbeck, B. B., & Romanski, L. M. (2006). Integration of auditory and visual communication information in the primate ventrolateral prefrontal cortex. Journal of Neuroscience, 26(43), 11138–11147.

Teng, S., Sommer, V. R., Pantazis, D., & Oliva, A. (2017). Hearing Scenes: A Neuromagnetic Signature of Auditory Source and Reverberant Space Separation. eNeuro, 4(1), ENEURO.0007-17.2017. http://doi.org/10.1523/ENEURO.0007-17.2017

Torralbo, A., Walther, D. B., Chai, B., Caddigan, E., Fei-Fei, L., & Beck, D. M. (2013). Good exemplars of natural scene categories elicit clearer patterns than bad exemplars but not greater BOLD activity. PloS one, 8(3), e58594.

Vetter, P., Smith, F. W., & Muckli, L. (2014). Decoding sound and imagery content in early visual cortex. Current Biology, 24(11), 1256–1262. http://doi.org/10.1016/j.cub.2014.04.020

Walther, D. B., Caddigan, E., Fei-Fei, L., & Beck, D. M. (2009). Natural Scene Categories Revealed in Distributed Patterns of Activity in the Human Brain. Journal of Neuroscience, 29(34), 10573–10581.

Walther, D. B., Chai, B., Caddigan, E., Beck, D. M., & Fei-Fei, L. (2011). Simple line drawings suffice for functional MRI decoding of natural scene categories. Proceedings of the National Academy of Sciences, 108(23), 9661–9666.

Walther, D. B., Beck, D. M., & Fei-Fei, L. (2012). To err is human: Correlating fMRI decoding and behavioral errors to probe the neural representation of natural scene categories. Visual population codes-Toward a common multivariate framework for cell recording and functional imaging, 391–416.

Wang, D., & Brown, G. J. (2006). Computational auditory scene analysis: Principles, algorithms, and applications. Wiley-IEEE Press.

Westfall, P. H., & Young, S. S. (1993). Resampling-based multiple testing: Examples and methods for p-value adjustment (Vol. 279). John Wiley & Sons.

Wood, J. N., & Grafman, J. (2003). Human prefrontal cortex: processing and representational perspectives. Nature Reviews Neuroscience, 4(2), 139–147. http://doi.org/10.1038/nrn1033.

